# The interplay between DNA methylation and sequence divergence in recent human evolution

**DOI:** 10.1101/015966

**Authors:** Irene Hernando-Herraez, Holger Heyn, Marcos Fernandez-Callejo, Enrique Vidal, Hugo Fernandez-Bellon, Javier Prado-Martinez, Andrew J Sharp, Manel Esteller, Tomas Marques-Bonet

## Abstract

DNA methylation is a key regulatory mechanism in mammalian genomes. Despite the increasing knowledge about this epigenetic modification, the understanding of human epigenome evolution is in its infancy. We used whole genome bisulfite sequencing to study DNA methylation and nucleotide divergence between human and great apes. We identified 360 and 210 differentially hypo- and hypermethylated regions (DMRs) in humans compared to non-human primates and estimated that 20% and 36% of these regions, respectively, were detectable throughout several human tissues. Human DMRs were enriched for specific histone modifications and contrary to expectations, the majority were located distal to transcription start sites, highlighting the importance of regions outside the direct regulatory context. We also found a significant excess of endogenous retrovirus elements in human-specific hypomethylated regions suggesting their association with local epigenetic changes.

We also reported for the first time a close interplay between inter-species genetic and epigenetic variation in regions of incomplete lineage sorting, transcription factor binding sites and human differentially hypermethylated regions. Specifically, we observed an excess of human-specific substitutions in transcription factor binding sites located within human DMRs, suggesting that alteration of regulatory motifs underlies some human-specific methylation patterns. We also found that the acquisition of DNA hypermethylation in the human lineage is frequently coupled with a rapid evolution at nucleotide level in the neighborhood of these CpG sites. Taken together, our results reveal new insights into the mechanistic basis of human-specific DNA methylation patterns and the interpretation of inter-species non-coding variation.

**Author summary:** Human and great apes, their closest living relatives, differ in numerous morphological and cognitive aspects, however their protein sequences are highly similar. It has long been hypothesized that human specific traits may be explained by changes in regulatory elements rather than by changes in primary sequence. In this context, evolutionary biologists have identified regulatory regions based on nucleotide sequence acceleration or conservation. However, epigenetics adds an extra layer of information that cannot be detected when comparing DNA sequences. The current study provides one of the first genome-wide comparison of genetic and epigenetic variation among humans and our closest living relatives. We identify and describe hundreds of regions presenting a human-specific DNA methylation pattern compared to great apes. We also report a local interplay between DNA methylation changes and the underlying nucleotide sequence in regions of incomplete lineage sorting and in regions of transcription factor binding, suggesting that both phenomena are closely associated.

## Introduction

A major aim of molecular biology is to understand the mechanisms that drive specific phenotypes. Humans and great apes differ in numerous morphological and cognitive aspects. However, their coding sequences are highly similar and most of the differences are located in non-coding regions [1], making it a challenge to define clear genotype-phenotype associations. It has been proposed that human specific traits originate from gene regulatory differences rather than from changes in the primary genetic sequence [2]. The characterization of regulatory domains is therefore a promising strategy to unveil regions of relevance for human evolution and to understand the implications of non-coding variation.

DNA methylation is a key regulatory mechanism of the genome [3]. It is present in many taxa and, in mammals, it plays an essential role in numerous biological processes ranging from cell differentiation to susceptibility to complex diseases [4, 5]. From a mechanistic perspective, DNA methylation has been described as an intermediate regulatory event, mediating the effect of genetic variability on phenotype formation [6]. However, the mechanisms by which the DNA methylation profile is generated are poorly understood. DNA methylation function is highly dependent on its location. In promoters, for example, it tends to confer gene repression while in gene bodies it is associated with transcriptional activation [3, 7]. DNA methylation levels also depend on the underlying genetic sequence and the occupancy of DNA binding factors [8, 9]. There is therefore no generic rule that can be applied to all biological situations, indicating the high complexity of the DNA methylation regulatory network.

In recent years, due to the development of genome-wide techniques that allow us to analyze DNA methylation profiles in multiple organisms, the field of comparative epigenomics has started to emerge. Exciting questions about how DNA methylation patterns vary through time and how this variation is linked to genome evolution can now be addressed. It has been shown that the global pattern of DNA methylation between close species, such as human and chimpanzee, is similar [10]. Nonetheless, there is a special interest in the study of local changes as mechanisms of species evolution, specially of human evolution [11]. Previous studies have identified several differentially methylated regions between human and primates using different techniques [12–19]. Interestingly, many of these regions have been associated not only with tissue-specific functions, but also with developmental and neurological mechanisms [13, 15, 19]. A key question that arises is how the epigenetic variability is transmitted across generations. The best studied mechanism is the dependence of DNA methylation levels on the genetic sequence. A growing number of studies in humans have shown an association between a nucleotide variant and a state of methylation [6, 20]. However the relationship between the genetic and the epigenetic sequence has not been explored when studying different species and despite recent advances in the field, many unanswered questions remain: How do DNA methylation patterns diverge across different genomic features? What are the processes driving such differences? Is this epigenetic variation associated with a higher rate of nucleotide substitution?

To further investigate these questions, we determined blood DNA methylation patterns in human, chimpanzee, gorilla and orangutan samples using whole genome bisulfite sequencing. Because this technique is not dependent on predefined sequences or methylation-dependent restriction enzymes, it is superior to other assays in analyzing patterns of DNA methylation [10, 12, 13]. We identified hundreds of human regions that differ in the DNA methylation pattern compared to the rest of great apes. These regions were enriched for specific histone modifications and they were located distal to transcription start sites. Furthermore, we found that DNA methylation variation and the underlying genetic code show close physical dependencies.

## Results

We performed whole genome bisulfite sequencing of blood derived DNA from a human (*Homo sapiens*), a chimpanzee (*Pan troglodytes*), a western gorilla (*Gorilla gorilla*) and a Sumatran orangutan (*Pongo abelii*) individual. A total of ~ 1.6 billion 100 bp Illumina paired-end reads were uniquely aligned to their respective reference genomes (hg19, panTro4 [1], gorGor3 [21], ponAbe2 [22]) using Bismark [23]. To facilitate an unbiased comparison between the four species, we performed all inter-species comparisons based on 6-primate EPO [24] restricting our analysis to 8,952,000 CpG sites conserved between the four species (see Methods). Compared to previous studies, this approach allowed us to reliably analyze a greater proportion of the species epigenomes. The read coverage in this subset of CpG sites averaged 10X in human, 12X in chimpanzee, 12X in gorilla and 13X in orangutan (Figure S1).

## A global view of Great Ape methylomes

Overall, the four species exhibited similar levels of DNA methylation with average levels of 74% in human, 71% in chimpanzee, 71% gorilla and 70 *%* in orangutan samples (Figure 1A). These findings are comparable to levels reported in previous studies analyzing blood methylomes [25, 26]. To investigate the epigenetic divergence between species, we retained 5,946,947 CpG sites that had between 4X-30X coverage in all species and performed correlation analysis in different regions of the genome. Here, the correlation coefficients of DNA methylation levels between species were in agreement with species phylogeny, the highest being in human-chimpanzee comparisons and the lowest in all comparisons involving orangutan (Figure 1B). From a genomic perspective, DNA methylation values correlated notably in promoters and CpG island regions and to a lesser extent at repeat loci (Figure 1B). Among the major repeat families, *Alu* elements presented the lowest correlation coefficients between species (Figure 1B). Only unique reads mapping to orthologous regions were considered and no major differences in coverage were observed among genomic locations (Figure S2 and Figure S3). However, due to the high frequency of CpG deamination in *Alu* elements, further studies are necessary to confirm the significance of this finding.

**Figure 1.**
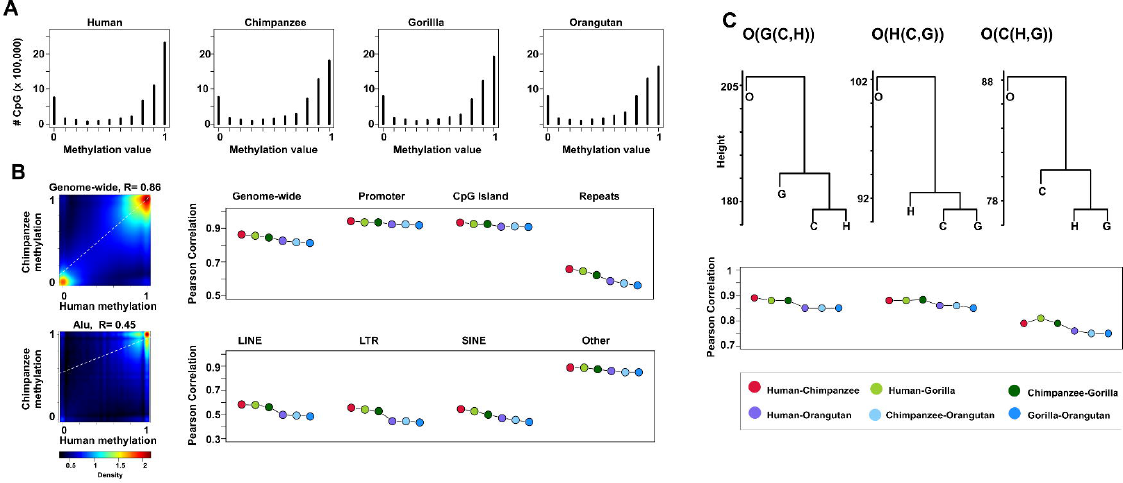
Global DNA methylation patterns. (A) DNA methylation profile of 5,946,947 CpG sites shared among the four species. (B) Pairwise-correlation analysis in different regions of the genome (right). Genome-wide n=5,946,947, promoter n=1,466,948, CpG Island n=740,153, repeats n= 2,310,842, LINE n=433,317, LTR n= 385,009, SINE 141,380, *Alu* 1,160,930, other n = 190,206. Density scatterplot of DNA methylation levels between human and chimpanzee genome-wide and in *Alu* elements (left), R indicates the Pearson correlation coefficient. (C) Hierarchical cluster tree and pairwise-correlation analysis based of methylation data from incomplete lineage sorting regions. O(G(C,H) n=922,701, O(H(C,G)) n= 221,908, O(C(H,G)) n= 142,231.

To gain insights into the relationship between nucleotide sequence and DNA methylation levels, we analyzed the DNA methylation patterns of regions whose sequence genealogy differs from the species phylogeny, known as regions of incomplete lineage sorting (ILS) [21]. Specifically, we studied regions where humans are more closely related to gorillas than to chimpanzees, represented as ((H,G)C)O) and regions where chimpanzees are more closely related to gorillas than to humans, represented as ((C,G)H)O). These regions are typically small (average length 473 bp) and contained 364,139 CpG sites conserved among all four species (Figure S4). Interestingly, hierarchical clustering and correlation analyses showed incomplete lineage sorting also at DNA methylation levels (Figure 1C), indicating a physical interplay between the genetic and epigenetic code and likely an epigenetic dependence on genetic mutations. While environmental heterogeneity can contribute to epigenetic variation [27] and therefore, it is a common confounding factor when comparing epigenomes of different species, the study of ILS regions allowed us to overcome this limitation. In addition, although 80% of the analyzed regions are not located either at promoters or within coding sequences, it has been shown that nearby genes present higher expression divergence between human and chimpanzee [21]. Consequently, we hypothesize that DNA methylation acts as a mediator mechanism, executing evolutionary driven commands of the genetic code and being involved in gene expression differences between species.

## Species-Specific DNA Methylation Patterns

We then focused our study on species-specific regions, which present a DNA methylation pattern exclusive to a single species. Therefore, we first identified hypomethylated regions (HMRs) throughout the genomes using a Hidden Markov Model [15]. This algorithm has been previously applied on human and chimpanzee DNA methylomes to detect putative regulatory regions [15, 28, 29]. We identified 28,835 (34.9 Mb) HMRs in human, 29,257 (33.6 Mb) in chimpanzee, 30,782 (36.5 Mb) in gorilla and 27,349 (33.1 Mb) regions in orangutan DNA methylomes. These hypomethylated regions were similar in size and methylation levels in all species (Figure S5) and harbored 15% of the CpG sites tested. Interestingly, an average of 72% (24.9 Mb) of HMRs were shared among all species and were mostly located in or close to human CpG islands (42.6% in CpG islands and 52.6% in CpG shores).

The resulting hypomethylated blocks were used to perform DNA methylation inter-species comparisons (see Methods). Due to the epigenomic differences between blood cell types [30], we required a stringent threshold of > 0.3 in mean CpG methylation difference (at least 30% methylation difference) to define a species-specific differentially methylated region (DMR) (see Methods). Moreover, this threshold allowed us to identify potential variant regions with higher biological impact. We defined two categories of DMRs: hypomethylated regions (in which one species is uniquely hypomethylated) (Table S2) and hypermethylated regions (in which one species is uniquely hypermethylated) (Table S3). Overall, we identified 360 hypomethylated DMRs in human (1.2% of HMRs), 340 in chimpanzee (1.1% of HMRs), 845 in gorilla (2.7% of HMRs) and 1,015 in orangutans (4.2% of HMRs) (Figure 2A). Further, we determined 210 DMRs specifically hypermethylated in human, 124 in chimpanzee, 167 in gorilla and 698 in orangutans (Figure 2A). One limitation of the method when calling a species hypermethylated DMRs is the intersection of multiple HMRs (from the other species) what resulted in smaller and fewer hypermethylated DMRs. Interestingly, species-specific hypomethylated regions were smaller in size (Wilcoxon test; P < 0.01) and had lower CpG density (Wilcoxon test; p<0.01) compared to HMRs common to the analyzed species. Due to the methodological limitations when calling hypermethylated DMRs, we could not assess size differences in this data set.

**Figure 2.**
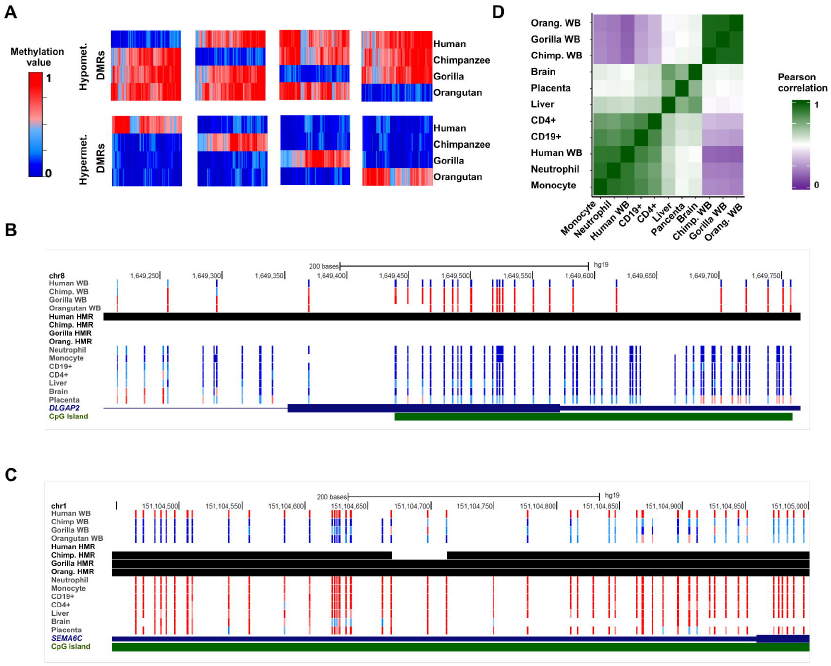
Differentially methylated regions. (A) Heat maps showing species specific hypo- (top) and hypermethylated (bottom) DRMs. Each vertical line represents the mean methylation value of a region. (B) Browser representation of human hypomethylated DMR,within DLGAP2 gene and (C) human hypermethylated DMR, within *SEMA6C*. Each vertical bar shows the methylation value of a single CpG site. Black blocks correspond to hypomethylated regions (HMRs) called by the Hidden Markov Model algorithm. Human samples: WB (whole blood), monocyte and neutrophil (myeloid lineage), CD19+ and CD4+ (lymphoid lineage), liver, brain and placenta. Non-human samples: WB: whole blood. (D) Pearson correlation matrix of human hypo- and hypermethylated DMRs.

An interesting example is represented by the last exon of the *DLGAP2* gene (Figure 2B), a membrane-associated guanylate kinase involved in synapse organization and signaling in neuronal cell [31]. This region was specifically hypomethylated in human whole blood compared to chimpanzee, gorilla and orangutan. Interestingly, we also observed hypomethylation at this region when comparing with published human methylomes (lymphoid [25] and myeloid cell types, and three other solid tissue types: brain, placenta and liver [32]]), indicating that this pattern is independent of cell types and likely to be conserved during development. An illustrative example of a human specific hypermethylated DMR which is conserved across both human hematopoietic cell types and solid tissues, is represented by the last exon and 3’UTR of the *SEMA6C* gene (Figure 2C). This gene encodes a member of the semaphorin family involved in axonal growth and synaptic connectivity maintenance [33]. Overall, we determined a strong correlation in the DNA methylation profile of human DMRs between the human whole blood sample and major hematopoietic cell types (Pearson’s correlation test, r^2^ > 0.8; Figure 2D). Herein, 66% of human hypomethylated DMRs and 64% of human hypermethylated DMRs were also detectable in the sorted blood cell types (mean difference < 0.3 between human whole blood and all cell types). In addition, 20% and 36% of human hypo- and hypermethylated DMRs, respectively, were detectable in all human tissues including solid cell types (brain, placenta, liver) (Figure 2D).

## Genomic Divergence in Differentially Methylated Regions

We further investigated the relationship between epigenetic and genetic evolution. Specifically, we aimed to determine the association between changes in DNA methylation and changes in the underlying genetic sequence. Therefore, we used the EPO multi-alignments blocks [24] to calculate lineage specific nucleotide substitutions that occurred in human DMRs and at their flanking regions (see Methods). We observed that human hypermethylated DMRs accumulated nucleotide substitutions in the same branch where the DNA methylation change occurred, clearly suggesting the epigenetic evolution to be coupled with nucleotide changes in these regions (Wilcoxon test, P < 0.05, Figure 3A). Moreover, due to the fact that hypermethylated cytosines deaminate spontaneously, we hypothesized a decrease in the number of CpG sites in the hypermethylated species as result of C>T mutations. However, no significant differences in CpG density were observed between species (Figure 3B) and no increase at C>T mutation was observed when classifying the human specific substitutions (Figure 3C), demonstrating that the increase of nucleotide substitutions is not due to cytosine deamination. Surprisingly, we instead observed an increase in the frequency of C>G mutations within human hypermethylated DMRs. Previous studies has pointed to oxidative conditions as the cause of this type of mutation [34], however further studies are required to interpret our finding.

**Figure 3.**
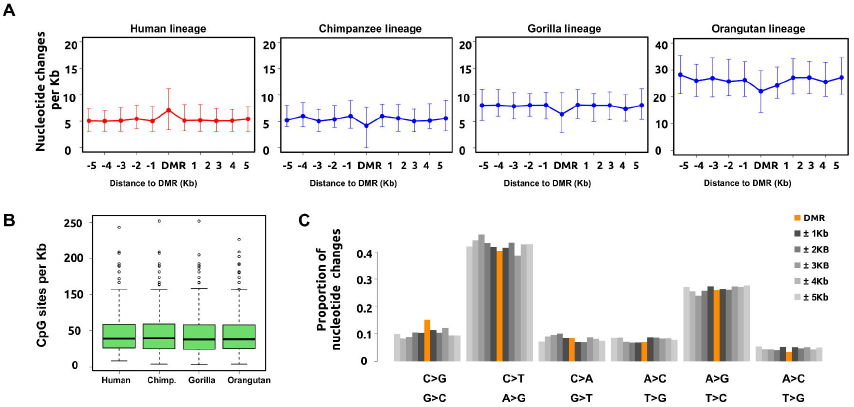
Nucleotide divergence at human hypermethylated DMRs. (A) Nucleotide changes of human hypermethylated DMRs estimated in each species lineage. The color plot represents the methylation state of the lineage species, red hypermethylated and blue hypomethylated. Data are represented as mean ± 2 standard deviations above and below the mean. (B) Number of CpG sites per Kb in human hypermethylated DMRs estimated at each species lineage. (C) Classification of human-specific substitutions showing an excess of C>G mutations at human hypermethylated DMRs compared to the flanking regions.

In contrast to the association of the genetic and epigenetic code in hypermethylated DMRs, human specific hypomethylated DMRs did not show significant differences in the rate of nucleotide substitutions compared to their flanking regions (Figure S6), suggesting that different mechanisms are implicated in the evolutionary gain and loss of CpG methylation.

## Functional Context of Human DMRs

To further investigate the mechanistic links between genetic and epigenetic changes we studied the DNA sequence of a high-confidence set of predicted transcription factor binding sites (TFBS) located within human-specific DMRs inferred using the CENTIPEDE algorithm [35]. We first identified 699 and 274 TFBS overlapping with hypo- and hypermethylated DMRs respectively, and compared these to 752,143 TFBS present in the 5,946,947 CpG sites data set (background). We then identified the proportion of binding sites whose DNA sequence is conserved between species and the proportion of binding sites containing at least one human specific change (Figure 4A). Within human-specific DMRs we observed a significant increase in the frequency of human-specific substitutions in TFBS when compared with TFBS in the background set (p < 0.001 hypomethylated DMRs and P = 0.008 hypermethylated DMRs, Figure S7), indicating a close evolutionary relationship between functional TFBS and local DNA methylation patterns.

**Figure 4.**
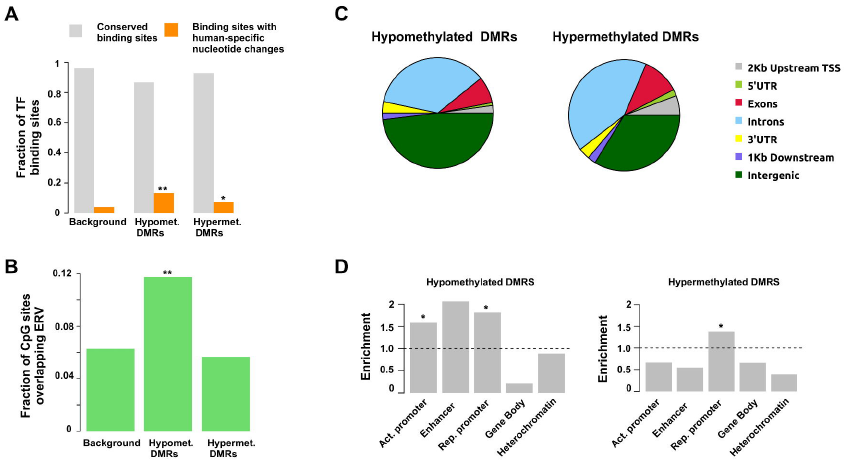
Characteristics of human DMRs. (A) Increase of human-specific substitutions in TFBS within DMRs compared with TFBS in the background set. Conserved binding sites (grey) and binding sites with human-specific changes (orange). (B) Fraction of CpG sites overlapping with ERV elements (C) Distribution of human hypo- and hypermethylated DMRs. (D) Histone modification enrichment at human hypo- and hyper DMRs. Active promoter: H3K9ac, enhancer: H3K4me1, repressive promoter: H3K27me3, gene body: H3K36me3 and heterochromatin: H3K9me3 **denotes p<0.001 and * denotes p<0.01 (permutation test).

We also examined the presence of common repetitive elements within human-specific DMRs. Interestingly, we found that 399 CpG sites located at human hypomethylated DMRs, representing 12% of all CpG sites located within hypomethylated human DMRs, overlapped with endogenous retroviruses (ERVs) (Figure 4B). This represents a two-fold enrichment over the background (P < 0.001, Figure S8). Recent studies indicate that ERV elements participate in transcriptional regulation during mammalian development [36]. Hence, our results suggest that ERV methylation levels could also be an important driving force in shaping primate epigenomes.

In total, 13.6% and 21.0% of hypo- and hypermethylated human DMRs, respectively, overlapped with promoter or exonic regions (Figure 4C). To investigate the functional role of the human DMRs, we determined their co-localization with histone mark occupancy data derived from chromatin immunoprecipitation sequencing (ChIP-seq) experiments in whole blood [37]. Specifically, we integrated DMRs with histone modifications data marking promoter, enhancer, gene body and heterochromatic regions (H3K9ac, H3K4me1, H3K27me3, H3K36me3 and H3K9me3). Here, we found that 52% of hypomethylated DMRs and 50% of hypermethylated DMRs co-localize with at least one histone modification. To determine a significant enrichment of a particular modification, we generated a set of random DMRs taking into account size, length, chromosomal location and CpG density. We observed that hypomethylated DMRs overlapped significantly with the actively regulating histone marks H3K9ac, H3K27me3 and H3K4me1, whereas hypermethylated DMRs were significantly enriched at loci marked by H3K27me3, regions with putative functions in developmental processes [38] (Permutation test P < 0.01, Figure 4D). In addition, we found that human DMRs were located distal to transcription start sites (TSS) compared to the distribution of the random DMRs (Figure S9). In particular, 44% and 37% of hypo- and hypermethylated DMRs, respectively, were located > 30kb away from the closest TSS (random DMRs: mean hypo =17%, mean hyper = 28%) (Permutation test P < 0.01 and P = 0.01, respectively). We therefore conclude that human DMRs are significantly enriched in regions occupied by active histone marks and are located outside the proximal gene regulatory context. Previous studies have shown that tissue regulatory events are mainly mediated by distal enhancers [26, 39]. Here, we propose enhancer and bivalent regions (H3K4me1 and H3K27me3) not only to be involved in the determination of cellular phenotypes, but also species phenotypes. However, further studies comparing DNA methylation maps from different tissues are required to understand tissue-specificity from an evolutionary point of view.

## Discussion

The current study provides one of the first genome-wide comparison of genetic and epigenetic variation among humans and our closest living relatives. Previous studies have analyzed a limited proportion of the genome using array methods or methylation-dependent restriction enzymes [6, 10, 12, 13, 17, 18]. Moreover the here applied use of EPO-alignments allowed us to cover a greater proportion of the epigenome in comparison to studies restricted to orthologous genes [14].

We found an overall conservation of the DNA methylation profiles between species. In concordance with previous studies [13, 40, 41], our results showed high conservation of DNA methylation levels specifically at CpG islands and gene promoters. Nevertheless, we found 570 regions that presented an exclusive pattern of DNA methylation in humans and contrary to expectation, these tend to be located distally to transcription start sites. This fact has to be taken into account in future evolutionary research, as to date most studies focused on the role of promoter DNA methylation and gene silencing. In this context, it was described that differences in promoter methylation underlie only 12-18% of differences in gene expression levels between humans and chimpanzees [12]. However, in our genome-wide study we have observed that most human-specific changes occur outside gene promoters. Because distal regulatory elements may contribute to transcriptional activity we hypothesize that the proportion of differences in expression levels explained by DNA methylation may be higher when analyzing whole-genome data sets. In this sense, this epigenetic variation could underlie tissue differences between species. Nevertheless, we have shown that 20% and 36% of human hypo- and hypermethylated DMRs, respectively, were detectable throughout several human tissues, suggesting their conservation during development. This phenomenon could also explain the discrepancy between certain DMRs and tissue function (for example, DMRs associated to neuronal genes detected in blood, Figure 2B and 2C) and highlights the importance of developmental and cell differentiation processes in the generation of species-specific traits.

Further, we have shown a close physical relationship between the genetic and the epigenetic code in three different analyses. Firstly, we have shown ILS of DNA methylation levels in regions that do not follow the species tree, suggesting a dependence of DNA methylation state on the underlying genetic sequence. Secondly, we have determined that a substantial and significant proportion of transcription factor binding sites at human DMRs contain human-specific mutations. This suggests a mechanistic link between the modification of binding sites at the nucleotide level and alterations of DNA methylation [42]. Thirdly, we found that the acquisition of DNA hypermethylation in the human lineage is frequently coupled with a rapid evolution at nucleotide level in the neighborhood of these CpG sites. This would initially suggest a loss of functionality of these regions and the subsequent accumulation of mutations. However, the observation of an enrichment for specific histone modification contradicts this hypothesis and rather point to a complex regulatory mechanism between histone modifications, DNA methylation and the underlying genetic sequence. The relationship between the species genetic background and the epigenetic code identified here also indicates that in these regions DNA methylation changes are not generated by stochastic events, environmental factors or cell type composition. The genetic-epigenetic association also suggests that DNA methylation patterns at these regions are a fixed feature in the human epigenome and excludes potential bias due to our limited sample sized. Importantly, the interplay between DNA methylation and sequence divergence reveal new insights into the mechanistic basis of human-specific DNA methylation patterns and the interpretation of inter-species non-coding variation.

## Materials and methods

The bisulfite sequencing data discussed in this publication have been deposited in NCBI’s Gene Expression Omnibus and are accessible through GEO Series accession number GSEXXXXX. *(authors note: a GEO submission is currently in progress - a valid accession ID will be provided prior to publication)*

## Sample Collection

Human and non-human research has been approved by the ethical committee of the European Research Union. No living animal has been used and all great ape blood samples were taken during routine health checks. Human donors gave written informed consent to take part in the study. Human blood was obtained from healthy donors and CD4/19 positive cells (T/B-cells) separated using the CD4+/19+ cell Isolation kit II (Miltenyi Biotec) following the manufacturer’s instructions [25]. DNA was extracted using phenol:chloroform:isoamylalcohol (Sigma). DNA methylation data from additional blood cell and solid tissue types were obtained from the Blueprint data portal (http://dcc.blueprintepigenome.eu/#/md/data) and previous publications (GSE46698), respectively.

## Library preparation

We spiked genomic DNA (1 or 2 μg) with unmethylated λ DNA (5 ng of *λ* DNA per μg of genomic DNA) (Promega). We sheared DNA by sonication to 50−500 bp with a Covaris E220 and selected 150- to 300 bp fragments using AMPure XP beads (Agencourt Bioscience Corp.). We constructed genomic DNA libraries using the TruSeq Sample Preparation kit (Illumina Inc.) following Illumina’s standard protocol. After adaptor ligation, we treated DNA with sodium bisulfite using the EpiTect Bisulfite kit (Qiagen) following the manufacturer’s instructions for formalin-fixed and paraffin-embedded (FFPE) tissue samples. We performed two rounds of conversion to achieve >99% conversion. We enriched adaptor-ligated DNA through seven cycles of PCR using the PfuTurboCx Hotstart DNA polymerase (Stratagene). We monitored library quality using the Agilent 2100 BioAnalyzer (Agilent) and determined the concentration of viable sequencing fragments (molecules carrying adapters at both extremities) by quantitative PCR using the Library Quantification Kit from KAPA Biosystems. We performed paired-end DNA sequencing (two reads of 100 bp each) using the Illumina Hi-Seq 2000. Sequencing quality was assessed using the Illumina Sequencing Analysis Viewer and FastQC software. We ensured the raw reads used in subsequent analyses were within the standard parameters set by the Illumina protocol. Positional quality along the reads was confirmed to be QC>30, and we excluded biases towards specific motifs or GC-enriched regions in the PCR amplification or hybridization.

## Mapping and Annotation

Paired-end sequencing reads (100 bp) were mapped to the *in silico* bisulfite-converted human (hg19), chimpanzee (panTro4) [1], gorilla (gorGor3) [21] and orangutan (ponAbe2) [22] references genomes using Bismark v0.7.8 [23] not allowing multiple alignments. We also removed potential PCR duplicates using Bismark’s deduplicate_bismark program. Custom Perl scripts were used to summarize the methylation levels of individual cytosines based on frequency of mapped reads.

To facilitate an unbiased comparison of the four genomes we used the Enredo-Pecan-Orthus (EPO) whole-genome multiple alignments of human, chimpanzee, gorilla, and orangutan [Ensemble Compara.6_primates_EPO] [24]. We identified 8,952,000 CpG positions shared among the four species in autosomal chromosomes, this data set was used for further analysis.

## Global methylome analysis

We used 5,946,947 CpG sites presenting a read coverage between 4X-30X in all species to perform global methylome comparisons according to their genomic annotation. Promoter regions were defined as +/- 2 Kb interval of the transcription start site. CpG island and repeat families were annotated using human UCSC Genome Browser tracks [43]. We used incomplete lineage sorting coordinates previously described [21].

## Identification of differentially methylated regions

Methylation values and number of reads in each position were used to identify hypomethylated regions (HMRs) using each reference genome coordinates by using a two-state Hidden Markov model [15]. The algorithm was developed to assess the methylation profile in humans and chimpanzees by dividing the methylome into regions of hypermethylation and hypomethylation. Non-human HMRs coordinates were converted hg19 coordinates using the EPO alignments. To call hypomethylated DMRs we first intersected a species HMRs with the other three methylomes and performed inter-species comparisons. To call hypermethylated DMRs, we intersected three species HMRs and then compared the methylation patterns to the methylome of the species of interest.

In order to define a species-specific differentially methylated region (DMR), we required a stringent threshold of > 0.3 in mean CpG methylation difference and a minimum of 5 CpGs (coverage between 4X-30X) in all species. Since methylation values can be interpreted as the percentage of methylation at a given site, a difference of 0.3 in CpG methylation indicates that there has been a change in methylation in 30% of the molecules tested. The proportion of cells present in blood, which are predominately neutrophils and lymphocytes, has similar proportions in chimpanzee, gorilla and orangutan [44, 45] (Table S4) and because our analysis required a mean methylation difference > 0.3 to be called DMR, changes in blood cell fractions representing < 30% of whole blood will unlikely affect our results.

## Genomic divergence and TFBS

We computed lineage specific nucleotide substitutions by extracting EPO multi-alignments blocks of human DMRs and flanking regions. Flanking regions were chosen with length equal to DMRs and located from 1 to 5 Kb upstream and downstream of DMRs. We then calculated the number of lineage specific nucleotides and divided by the amount of nucleotides present in the four species. Insertions and deletions were not taken into account in this analysis. Transcription factor binding sites coordinates were previously identified [35] and human specific substitutions were also calculated using EPO multi-alignments blocks.

## Histone modification enrichment

The genomic distribution shown in Figure 4A, was performed considering the human hg19 RefGene annotation using PAVIS [46]. We used processed ChIP-seq data previously published [37]. To determine enrichment and significance of a particular modification, we generated 100 control sets sized-matched of the human hypo- and hypermethylated DMRs independently. To generate this control data set we also took into account chromosome location, CpG density and length. Next, we determined the proportion of each histone codification overlapping the human DMRs and the control data sets. The ratio of the two is reported as enrichment shown in Figure 4D. The P-value corresponded to the number of times that the DMRs proportions appeared in control data set distribution, divided by the number of sets (n = 100). Similarly, to determine the significance of DMRs location we calculated the proportion of DMRs ± 30 Kb around TSS (RefSeq genes) and compared to the control data set distribution. The p-value corresponded to the number of times that the DMRs proportion appeared in control data set distribution, divided by the number of sets (n = 100) (Figure S9).

## Acknowledgments

We gratefully acknowledge Teresa Abello from Barcelona Zoo and Christina Hvilsom from Copenhagen Zoo for providing non-human samples.

## Supporting information

**Figure S1:** Read coverage across samples.

**Figure S2:** Read coverage across genomic regions and samples.

**Figure S3:** Read coverage across major repeat families and samples.

**Figure S4:** Size of incomplete lineage sorting regions.

**Figure S5:** Distribution of (A) size and (B) methylation value of hypomethyated regions (HMRs) across samples.

**Figure S6:** Nucleotide changes of human hypomethylated DMRs estimated in each species lineage. The color plot represents the methylation state of the lineage species, red hypermethylated and blue hypomethylated. Data are represented as mean ± 2 standard deviations above and below the mean.

**Figure S7:** Proportion of binding sites with human-specific nucleotide changes at DMRs (green) and background (black). (A) Human hypomethylated DMRs. (B) Human hypermethylated DMRs.

**Figure S8:** CpG sites within a ERV repeat elements at human hypomethylated DMRs and background.

**Figure S9:** (A) Distance distribution of human hypo- and hypermethylated DMRs to the closest RefSeq TSS truncated at 100kb. (B) Proportion of human DMRs (red) and control datasets (black) located > 30kb away from the closest TSS.

**Table S1:** Species specific hypomethylated DMRs.

**Table S2:** Species specific hypermethylated DMRs.

**Table S3:** Proportion (%) of neutrophils and lymphocytes in whole blood.

